# Empirical evidence that diversionary feeding increases productivity in ground-nesting birds

**DOI:** 10.1101/2024.12.06.627135

**Authors:** Jack A. Bamber, Kenny Kortland, Chris Sutherland, Xavier Lambin

**Author notes:** **Author Contributions** *Conceptualisation*: Kortland, Bamber, Lambin, Sutherland; *Fieldwork*: Bamber; *Statistical analysis*: Bamber, Lambin, Sutherland; *Data Curation*: Bamber; *Writing- original draft preparation*: Bamber; *Writing – review and editing*: Bamber, Sutherland, Kortland, Lambin; *Funding acquisition*: Lambin, Kortland, Sutherland. *All authors have read and agreed to the published version of the manuscript*.

## Abstract

The recovery of predator populations can negatively impact other species of conservation concern, leading to conservation conflicts. Evidence-based solutions are needed to resolve such conflicts without sacrificing hard-won gains for recovering species. Well-designed, large-scale field experiments provide the most rigorous evidence to justify new forms of intervention, but they are notoriously hard to implement. Further, monitoring scarce species without negative impacts is challenging, calling for indirect and non-invasive monitoring methods. Uncertainties remain about whether observational monitoring adequately reflects the true processes of interest.

Having conducted a well-designed, large-scale, diversionary feeding field experiment that reduced artificial nest depredation, we evaluated whether this translated to capercaillie productivity in the same area. Using camera traps aimed at dust baths, we non-invasively monitored capercaillie hen productivity over 3 years and in 30 1km^2^ grid cells under a randomised control (un-fed) and treatment (fed) design. Diversionary feeding significantly increased the probability that a detected hen would have a brood. The impact of diversionary feeding did not change over the brooding season, indicating that hens without a brood had failed due to nest depredation rather than predation of chicks. The probability of detecting a hen with a brood was 0.85 (0.65-0.94) in fed locations, more than double that in unfed locations, which was 0.37 (CI 0.2-0.57). The average brood size was reduced over time, but the change did not differ between fed and unfed sites. This is in line with natural mortality occurring independently of diversionary feeding. Importantly, the increased chance of having a brood in the fed areas and the predicted brood size leads to a substantial increase in overall productivity – the expected number of chicks per hen – at the end of the sampling season. This was just 0.82 (0.35 – 1.29) chicks per hen in the unfed sites and more than double 1.90 (1.24 – 2.55) chicks in fed sites.

This study provides compelling empirical evidence that diversionary feeding positively affects productivity, offering an effective non-lethal solution to the increasingly common conservation conflict where both predator and prey are afforded protection.

## Introduction

Robust evaluation of management strategies is vital to support decision-making [1]. However, some historically successful management interventions have become less suitable in the contemporary context, including where species assemblages are reforming under legal protection and conservation efforts [2,3]. Despite this, many traditional management interventions are still used despite little evidence of their suitability and, in some instances, despite evidence they are ineffective under current conditions.

Lethal control of predators to maintain a harvestable surplus of prey species and to protect livestock remains a widely used management practice [4]. In its current form, lethal predator control delivers only short-term reductions in abundance owing to compensatory immigration and breeding and, therefore, requires repeated (i.e., annual) implementation [5,6]. Protection of some predator species also reduces the efficacy of lethal control because changes in predation rates by the non-controlled predators can also be compensatory [7]. Moreover, social norms increasingly challenge the legitimacy of lethal control, whether for the maintenance of game species or the protection of species of conservation concern [8]. The lack of long-term efficacy, moral objections, and the differing management practices across landownerships contribute to a growing pressure to find alternatives to lethal control. Yet, barriers exist to the uptake of alternatives, such as the attachment to traditional practices and a lack of reliable or trusted evidence on their efficacy and practicality [9].

Controversies around predator control have become acute in the context of recovering predators, especially when it results in increased predation by recovering predators on declining or valued prey species [10,11]. For example, predation by meso-carnivores on nests and chicks of ground-nesting birds is a key driver in the decline of ground-nesting bird populations [12,13]. In such circumstances, relieving predation pressure on prey populations may be necessary to restore productivity and allow prey population persistence [14]. However, there remains uncertainty about the circumstances under which non-lethal interventions offer a feasible solution to these conflicts.

One promising non-lethal intervention to reduce predation impact is diversionary feeding, the deliberate provision of alternative food to divert predation pressure away from species of management interest [15,16]. It is predicated on the premise that predators will preferentially consume readily accessible, provisioned food rather than searching for cryptic nests and chicks [17]. Diversionary feeding has been demonstrated to reduce predation significantly in both game and threatened bird species by various predators [18–20]. However, the extent to which the foraging behaviour of problematic predators can be modified through management is unknown in most specific contexts. Accordingly, strong evidence for the effects of diversionary feeding is necessary before it is deployed as an alternative to traditional lethal control.

In most jurisdictions, legal protection of predators prevents population control unless there is evidence that culling predators under licence would achieve specific management objectives. This is another area of uncertainty, as direct evidence is rarely available. This increases the polarisation between proponents of traditional lethal control and conservationists unwilling to sacrifice recent conservation gains with the resumption of culling [11].

The conservation conflict between the legally protected and recovering pine marten (*Martes martes*) and the drastically declining capercaillie (*Tetrao urogallus*) is a good example of a context where diversionary feeding holds promise. Pine martens are seasonal nest predators implicated in the continued decline of ground-nesting capercaillie [21,22]. A randomised experiment recently demonstrated an 83% reduction in depredation rates on artificial nests by pine marten with short-term diversionary feeding [16]. Using artificial nests, as they did, as a proxy for nest predation allowed experimental rigour but fell short of providing direct evidence that diversionary feeding does reduce predation on actual capercaillie nests. A significant evidence gap remains in our understanding of the beneficial impact of diversionary feeding for target species. In this study, we directly address this knowledge gap using a landscape-scale feeding experiment to quantify the effects of diversionary feeding on capercaillie productivity – measured as the expected number of chicks per hen. Through non-invasive monitoring of capercaillie broods over time, we quantify to what extent diversionary feeding reduces the impact of predation on capercaillie breeding success.

## Methods

### Study Area

This study was conducted within the Cairngorms Connect landscape, a 600km^2^ multi-stakeholder ecological restoration project in the Cairngorms National Park, Scotland (57°09’47.5”N 3°42’47.0” W). The area is within the stronghold of Scotland’s largest remaining capercaillie populations [23] and hosts a recovering predator guild, including the pine marten. Importantly, no lethal predator control was conducted during sampling and was limited in the preceding 3 years. A vital aspect of managing the landscape is controlling red and roe deer (*Cervidae*) for forest regeneration. This provides both a year-round potential subsidy to predators and an easily accessible food source for diversionary feeding. Partner organisations sought non-lethal ways to manage predator impacts without resorting to population control. This experiment allowed the deployment and assessment of diversionary feeding as an emergency intervention, with the option to halt if adverse effects were observed.

### Diversionary feeding experimental design

The experiment was performed in a control (unfed) and treatment (fed) design. Unfed and fed sampling grid cells were deployed randomly across the experiment (Bamber *et al*. 2024). Sites used in this study were a subset of grid cells with recent records of capercaillie presence (historical brood counts, capercaillie sightings and lek activity from 2018 onwards, to provide an increased chance of capercaillie detection.

Grid cells were 1km^2^ in size, separated by a 1km^2^ spacing cell to maximise independence of the treatment. The size of sampling units was chosen to encompass the typical daily movement range of a pine marten, the focal nest predator (49 ha in females and 54 ha in males [24]). This size also encompassed the average home range of capercaillie broods, 0.4km^2^ [25], making it unlikely that broods would use multiple grid cells during experimentation. The fed grid was switched each year across three years of sampling (2021-23) to reduce the influence of cell properties that were not accounted for through the initial randomisation of grid cell selection.

Diversionary feeding stations were deployed in 2021, 2022, and 2023, from the last week of April to the first week in July, encompassing capercaillie egg laying, incubation and the first 30 days after hatching. Approximately 10kg of deer carrion were placed at each feeding station and restocked every two weeks to ensure fresh food was always present. Any meat remaining when restocking was left in place to build scent cues to increase detection by local predators. Stations were deployed within ∼100m of the centre of the fed grids, avoiding water courses by >20m and human tracks and trails by >50m. The flipping of treatments between years meant that grid cells had the same treatment in 2021 and 2023, but feeding stations were at least 50m from the previous location. The short feeding timeframe and changing of feeding locations across years aimed to reduce the risk of increasing population size or causing long-term spatial aggregation of predators, respectively. For full details of the core experimental design, see Bamber et al (2024).

### Brood monitoring through camera trapping

To detect capercaillie non-invasively, camera traps (Browning Recon Force Advantage and Recon Force Elite; set at 3-shot burst with 5-second intervals) were deployed on dust baths (microsites known to be used by capercaillie broods (Bamber et al., 2023). Thirty-three grid cells were initially identified as having a historical capercaillie presence, making them suitable for cold search for camera deployments.

Cold searching consisted of a central loop within the sampling grid cell, scanning initially for capercaillie signs (feathers, droppings or sightings) to trigger localised searching of dusty areas (upturned root plates, exposed earth in hillsides, or bare earth in standing tree roots). Searching dusty areas triggered camera deployment if signs of use, dust bowls, capercaillie droppings, and capercaillie feathers were detected. Cameras were deployed in 30 of the 33 historical capercaillie grids with contemporary evidence of occupancy. If several dust baths were in proximity, baths with hen feathers and fresh signs of recent use were selected for deployment. If multiple high-quality dust baths were within a grid cell, dust baths farthest away from each other were selected. If no suitable dust bath with signs of use could be found, no camera was deployed. Cameras were deployed in clusters (2 to 4) within grid cells to increase the probability of detecting the focal species when present [27].

Monitoring of dust baths took place from late May to mid-September and included the most vulnerable period for recently hatched capercaillie, which hatch in early June, rely on their mothers in the first few weeks of life, and follow their mothers through to dispersal in early autumn.

### Data extraction

Camera trap images were processed primarily using the Digicam and CamTrapR framework to tag and extract detections of capercaillie, tagging males, females, and broods as three different types of detection [28]. Detections were separated by day for independence. The tagged images from 2021 and 2022 were used to train the Conservation AI software to automatically identify capercaillie cocks and hens to extract capercaillie images passively from 2023 [29]. We extracted encounters identified by conservation AI as containing capercaillie hens, and a manual assessment was performed on this subset of images using the same workflow as 2021 and 2022 to identify hen detections (removing false positives). Images were correctly identified to contain a capercaillie hen 95.4% of the time (13,532 correct, 14,119 identified). Brood counts were created manually from detections of capercaillie hens, assigning a chick count to each image during the encounter (all images detected in sequence from arrival at the dust bath till leaving). The chick count assigned to the individual encounter for analysis was the most chicks seen on an image in a sequence. This provided the response variable for modelling, the presence of a brood (0 or 1) and, if chicks were detected with a hen, the brood size (range: 1 – 8). Total detections per year and grid varied (See table 1), with extremes, one grid detecting a hen capercaillie on 29 separate days and others having only one detection across the entire sampling period.

**Table 1.** Camera deployments during sampling. Separated into individual cameras and Unique sites (independent sampling grid cells). The number of cameras and grid cells, alongside the number of sites detecting a capercaillie hen (with or without a brood) and a capercaillie brood, are presented.

| Year | Cameras |  |  |  | Unique sites |  |  |
| --- | --- | --- | --- | --- | --- | --- | --- |
|  | Deployed | Cameras with a detection | Hen detections | Hen with brood | Sites with cameras deployed | With hen detection | Hen with brood |
| 2021 | 57 | 24 | 47 | 23 | 25 | 16 | 12 |
| 2022 | 87 | 24 | 57 | 36 | 28 | 16 | 13 |
| 2023 | 102 | 60 | 204 | 122 | 30 | 25 | 16 |
| Total | 246 | 108 | 308 | 181 | 83 | 57 | 41 |

### Statistical analysis

Our inference objective was first to estimate the probability that a hen would have a brood (reproductive success) and second, conditional on a hen having a brood, the size of the brood (productivity). As such, we used a zero-truncated (or hurdle) Poisson regression model.

The zero-truncated model assumes that the zeros are binomial random variables, where barren = 0 and brood = 1, and was modelled using binomial regression with a logit link function. The number of chicks, conditional on a hen having a brood, was then assumed to be a truncated Poisson random variable modelled using a Poisson regression and a log link function. Importantly, the probability mass function of the Poisson model is truncated at 0 (i.e., does not include 0) and scaled accordingly. From this model, predictions can be made about the probability of having a brood, the expected brood size independent of having a brood and, importantly, the expected number of chicks conditional on having a brood. The latter provides an intuitive estimate for the number of chicks per hen, a standard measure of productivity for birds.

The binomial response and the conditional counts can be modelled as a function of covariates using standard generalised linear modelling approaches. Given the experimental nature of our study, we considered two covariates as predictors for both processes. We included a binary treatment effect (fed/unfed) to explicitly test for the effect of diversionary feeding on both the probability of a hen being barren and brood size. We also included a time of year effect (ordinal day) to account for the potential for brood presence and brood size to vary (i.e., decrease) over the 10-week study period. Ordinal day ran from day 0, relating to the 5^th^ of June, to day 98, relating to the 12^th^ of September, encompassing the period chicks are with hens. We included treatment-time interactions for both processes to test for treatment-specific temporal changes in the presence and number of chicks. To account for the non-independence of detections in the same sample grid (we deployed, on average, 3 cameras per grid), we used grid-year as a random effect, resulting in 57 grid-year sites in total.

Considering all combinations of covariates resulted in a total of 25 models. Models were fit in R (R Core Team) using the GLMMTMB function from the “glmmtmb” package [30]. Model selection was conducted based on AIC model comparisons [31] produced from the “glmmtmb” models, and predictions and 95% confidence intervals were calculated using the “ggeffects” package [32].

The estimates from the modelling described above provide information that can be used to parameterise a deterministic population projection model to explore the importance of diversionary feeding-induced changes in productivity on population growth. To do so, we adapt the parameters from the post-breeding capercaillie population projection model of Moss et al. (2000), which was developed for the same population. We parameterised the breeding probability using data from 13 tracked hens’ [33] productivity rates using our estimates of productivity (number of chicks per hen) from fed and unfed sites. Deterministic population growth rates and projections for populations with and without diversionary feeding were performed in R using the “popbio” package [34].

## Results

### Brood analysis

The presence or absence of diversionary feeding affected the probability that a hen was detected with a brood. However, this probability did not change over time. The top supported model, using AIC, included the effect of treatment only on the probability of having no brood, i.e., the binomial ‘zeros’ model, and the effect of ordinal day only on the expected brood size, i.e., the truncated Poisson count model (Table 2). This means that, specifically, diversionary feeding significantly reduced the probability of being barren from 0.848 (95% CI: 0.66 - 0.94) in unfed sites to 0.37 (0.57 - 0.94) in fed sites (estimate: -2.27, SE: 0.71, Figure 2). Note that the binomial model estimates the probability of a zero (barren), and one minus that probability is the probability of having a brood. We report this complement from here for clearer intuition (Figure 1).

**Table 2.** AIC table to show a model selection of variables; table contains the top 4 models with ΔAIC_c_ <2, out of the possible 25 model combinations, across the two parts of the hurdle model, Treatment and Ordinal day.

| Description | AIC | $\Delta AIC$ | Pars |
| --- | --- | --- | --- |
| Count: day of season<br>Zero: treatment | 983.311 | 0 | 4 |
| Count: day of season<br>Zero: treatment & day of season (interaction) | 984.614 | 1.303 | 6 |
| Count: day of season<br>Zero: treatment and day of season (additive) | 984.718 | 1.408 | 5 |
| Count: day of season & treatment (additive)<br>Zero: treatment | 985.137 | 1.826 | 5 |

**Figure 1.**
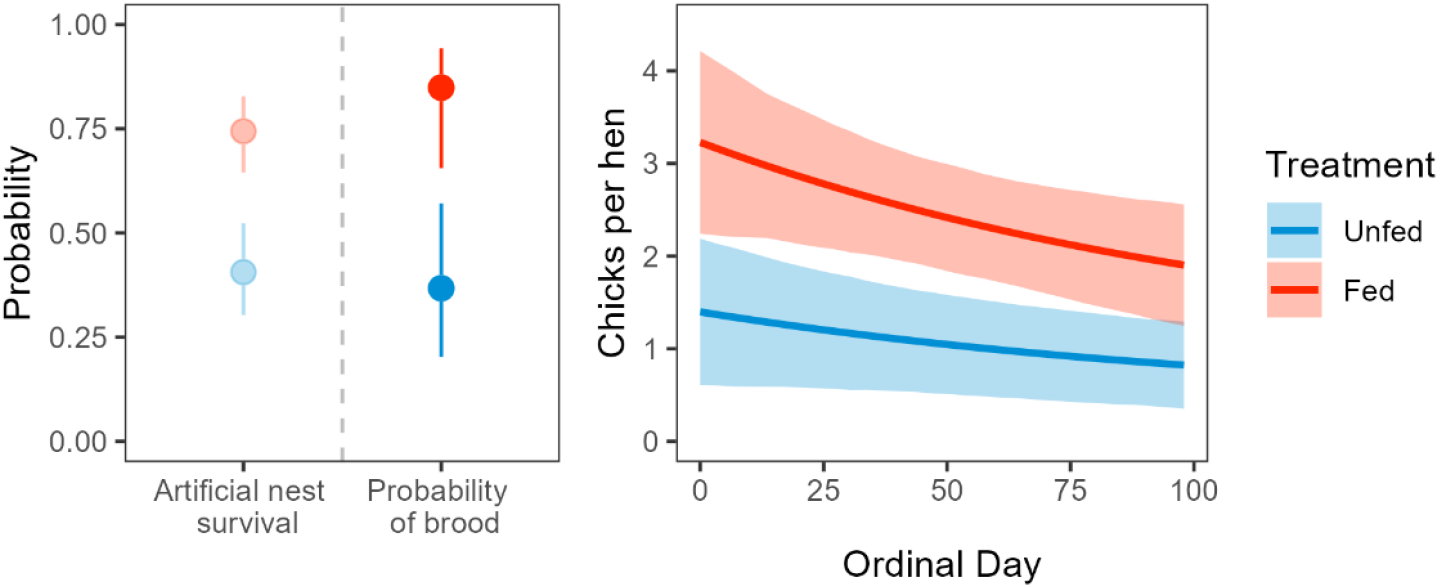
Impact of diversionary feeding on capercaillie productivity. Left panel: the probability of an artificial nest survival over 28 days from Bamber et al. (2024), the probability of detecting a hen with a brood. Right panel: the combined estimate of average chicks per hen over time, Day 0 (June 5^th^). The 95% confidence intervals are depicted in corresponding treatment colours: red for fed and blue for unfed.

In terms of inference about brood size, our results suggest that productivity declines throughout the study (−0.007, SE 0.002), but is not affected by the presence of diversionary feeding. Specifically, this means that, while the probability of having a brood is higher in fed sites, the expected brood size for a hen with a brood is not different and that the expected brood size declines over time at the same rate in fed and unfed sites (Figure 2). Importantly, the chicks per hen must include both the brood count model and the probability of having a brood, and this results in a stark productivity difference in fed versus unfed sites. For example, by the end of sampling (ordinal day 98), the predicted number of chicks per hen is 1.90 (1.24 – 2.55) in the fed sites, which is 131% higher than in the unfed sites, which is 0.82 (0.35 – 1.29, Figure 2).

**Table 2.** Coefficients of Zero Truncated hurdle model showing changes in the probability of a barren hen being detected (Zero) and the brood size change when a brood was detected (Count). Intercept for count is Day 0 (June 5^th^), for zero Treatment; Unfed). Dashed are included for effects not present in the count or zero sub-models.

|  | Count (Poisson, log link) |  |  | Zero (Binomial, logit link) |  |  |
| --- | --- | --- | --- | --- | --- | --- |
|  | Estimate | Std. Err. | p-value | Estimate | Std. Err. | p-value |
| Intercept | 1.311 | 0.124 | < 2e-16 | 0.543 | 0.423 | 0.199 |
| Treatment: Fed | - | - | - | -2.268 | 0.712 | 0.001 |
| Ordinal Day | -0.007 | 0.002 | 0.007 | - | - | - |

Combined model, accounting for the count combined with zero probability.

### Projected influence of productivity increase

Using our estimates of chicks per hen, we compared population growth rates under two feeding scenarios: 1) no-feeding and 2) implementation of diversionary feeding. In the unfed scenario, the growth rate is λ= 0.903, a population decline that is broadly consistent with predictions from Moss et al. (2000). The introduction of diversionary feeding into the model increases the population growth rate to λ=1.109. Thus, widescale adoption of diversionary feeding could boost population growth by 20%, an increase sufficient to reverse the predicted decline of capercaillie.

## Discussion

We found that diversionary feeding increases capercaillie productivity. In doing so, we demonstrated that the reduction in predation rates observed in a field experiment using artificial nests as a proxy response variable fully translates to an almost identical increase in the survival of actual capercaillie nests and, hence, subsequent productivity. Diversionary feeding increased the predicted number of chicks per hen detected at the end of the dependence period by over 131%, from 0.82 to 1.9 chicks per hen. While our result affirms that nest depredation significantly contributes to breeding failure in ground-nesting capercaillie [35], there is now a well-supported, non-lethal, feasible solution to this conservation conflict. Through strong inference afforded by a rare randomised, replicated field experiment, there arguably is now “certainty of the evidence” that diversionary feeding is demonstrably effective in reducing the impact of predation on capercaillie nesting success without undesirable harms as per the conservation evidence framework [36]. Accordingly, practitioners can now consider deploying this method with the knowledge that the barrier of lack of evidence has been addressed.

Bamber et al. (2024) found that nest survival increased to 74% with diversionary feeding, compared to 40% without it, mainly due to a significant decrease in depredation by pine martens and, to a lesser extent, badgers. Legitimate concerns remained that the use of artificial nests, including differences in predator behaviour between natural and artificial nests and the influence of scent cues (both increasing or decreasing predation), may have caused spurious results [37,38]. For instance, few artificial nests (1.3%) were consumed by crows (C*orvus corone*) and foxes (*Vulpes vulpes*), despite previous evidence of capercaillie nest depredation by these species [39,40] but mixed evidence on the actual rates of corvids [39]. The remarkable quantitative alignment between the probabilities of detecting a hen with a brood and of artificial nest survival (Figure 1) alleviates these concerns about using a proxy response variable in Bamber et al. (2024). Thus, diversionary feeding effectively reduces (mostly badger and pine marten) predation, obviating the need for lethal population control of these protected species. By implication, the quantitative contribution of crows and foxes must be small. Diversionary feeding is a viable tool to reconcile predator recovery with the seemingly conflicting conservation goals of protecting capercaillie.

Whether diversionary feeding reduced chick predation to the same extent as it reduced depredation of nests was a critical knowledge gap. We found no positive effect of our experiment on either brood size or its decay rate over time, indicating that diversionary feeding does not reduce chick predation. While predation likely contributes to chick loss, the guild of predators responsible is likely wider, and includes raptors. For example, goshawks, which predate capercaillie, and chicks [41,42], were only detected once across 75 diversionary feeding deployments. Therefore, experiment-induced reductions in predation by the predator species that did not utilise the provisioned food was not expected. Brood size declined significantly throughout the season, with predictions from the count model showing a decline of 51% (from 3.7 to 1.9 chicks) over 98 days. Outside predation, multiple factors have been linked to chick deaths in capercaillie, most notably cold and wet springs, which have been invoked as contributors to brood decay through their impact on insect prey availability [35,43]. As climate change progresses, these impacts may worsen, further reducing productivity. Diversionary feeding in this form is seemingly unsuitable if predator intervention aims to increase chick survival instead of the proportion of hens with broods.

Diversionary feeding more than doubled the predicted productivity to 1.9 chicks per hen. Diversionary feeding achieves this by increasing the number of broods rather than influencing brood size. This far surpasses the 0.6 to 1.1 chicks per hen previously suggested as necessary for population stabilisation in Scotland [44]. Projecting population growth with the increase in chicks per hen brought about by diversionary feeding shifted the annual population growth rate from a 9% decline to a 10 % increase. While there is more work to be done on integrating several factors that affect growth, this exercise helped demonstrate the potential and promise of diversionary feeding on the population growth of a species that many suggest is all but extinct. Our key message is that diversionary feeding can substantially mitigate the impact of nest predation on population growth, consistent with the prediction that nest loss more than chick loss impacts productivity [35].

### Wider Management Implications

Predicted chicks per hen more than doubled when the impact of predation was reduced by diversionary feeding. This improved productivity matches those seen in capercaillie populations at the top of the productivity range, often linked to peak vole years when depredation is deemed lowest due to abundant alternative prey (Appendix 1, [45]). Our evidence indicates that similarly high productivity rates could be realised by managing the impact of predation in other parts of the southern range of capercaillie (Cantabria, Alps, Bavaria), where predation is also deemed to contribute to rapid population declines [21,46]. Our evidence unambiguously shows that diversionary feeding effectively mitigates the impact of multiple predators, providing a tool for land managers wishing to intervene effectively on predator impact.

A barrier to action has been the concern raised by practitioners that feeding predators will increase overall predator abundance or cause local aggregation [15]. The feeding protocol we tested involved a short, 8-week-long seasonal feeding pulse that matched egg laying and incubation. This was short enough to minimise any impact on predator survival and to avoid a numerical response. The lack of a negative impact of diversionary feeding on brood size in this study may indicate that there is no numerical response associated with sites by predators. Notably, Olejarz & Podgórski (2024) found no consistent effect of supplementary feeding on the home ranges of terrestrial mammals, with feeding choices likely impacting at an individual level and not at a population level, implying that uniform aggregation is unlikely. Managers considering diversionary feeding should not let a fear of aggregation prevent an investigation into the method under their specific contexts.

Managing the impact of increased predation by generalist predators that have recovered from past persecution and now thrive in anthropogenic landscapes is a global issue in conservation [10,47]. Substantial resources are expended in managing predation to rescue an ever-growing list of at-risk, ground-nesting birds, such as godwits, common curlews, sage grouse, and black stilt, from declines [13,48–50]. In many cases, elevated predator abundances are symptoms of modified ecosystems rather than the ultimate cause of the decline, meaning that population control could negatively impact native predators without any long-term benefits to prey. Intervention to aid in slowing declines is desirable but should not come at the cost of broader conservation due to these conflicts [51]. Our evidence shows significant potential for impact-based intervention to alleviate unwanted predation and indicates the viability of coexistence conservation strategies within predator conflicts.

## Acknowledgements

We acknowledge funding from NERC SUPER DTP studentship to JB (NE/S007342/1) and Forestry and Land Scotland. We thank all Cairngorms Connect partner organisations, funders and staff for in-kind contributions, including but not limited to B.Innes, C.Watson, T.Cameron (FLS); A.Poulsen, T.MacDonell, D.McGibbon, D.Ross, R.Dugan, G.Ashley, M.Willis (Wildland Ltd); R.Mason, M.Butler, C.Waite, F.Cormack, J.Ward (RSPB); and the team at Conservation AI. Special thanks to R.Moss for sharing expertise on brood detections.

## Appendix 1

**Figure 4.**
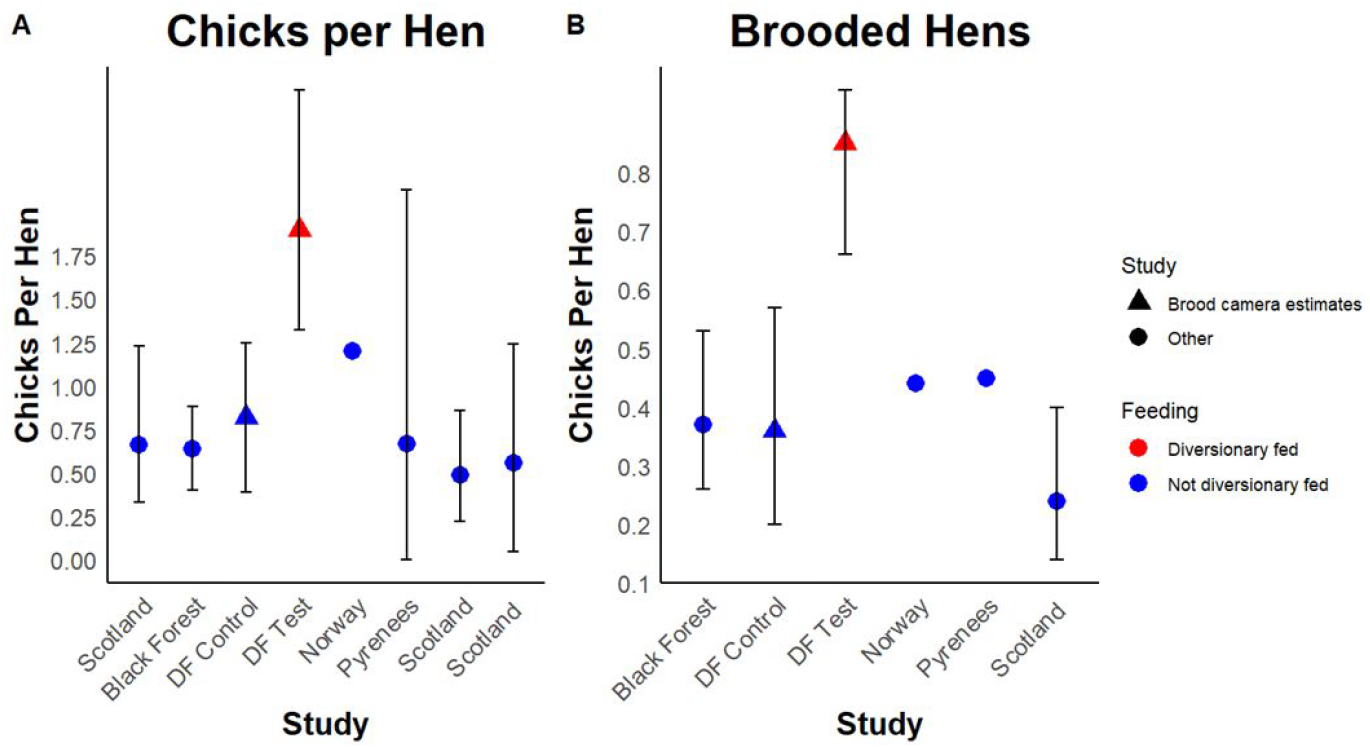
Collection of productivity estimates for capercaillie recruitment across Europe. A) shows chicks per hen, and B) shows brooded hens. Studies from the literature are shown in circles, with estimates from this study shown with triangles, and diversionary fed prediction is shown in red. When available, the range of estimates was included with black error bars (Baines & Aebischer, 2023; Coppes et al., 2021; Fletcher & Baines, 2020; Gil et al., 2020; Ruottinen et al., 2024; Summers et al., 2015; Wegge et al., 2022).

